# Quercetin prevents insulin dysfunction in hypertensive animals

**DOI:** 10.1101/2021.09.10.459841

**Authors:** Cristiane Alves Serra, Alexandre Freire dos Reis, Bruno Calsa, Cintia Sena Bueno, Júlia Venturini Helaehil, Suelen Aparecida Ribeiro de Souza, Camila Andrea de Oliveira, Emerielle Cristine Vanzella, Maria Esméria Corezola do Amaral

**Affiliations:** Graduate Program in Biomedical Sciences, Centro Universitário da Fundação Hermínio Ometto, FHO, Araras, SP, Brazil; Biomedical College, Centro Universitário da Fundação Hermínio Ometto, FHO, Araras, SP, Brazil; Departamento de Biologia Estrutural e Funcional, Instituto de Biologia, Universidade Estadual de Campinas, UNICAMP, Campinas, SP, Brasil

**Keywords:** hypertension, pancreatic islets, blood vessels, muscarinic receptors, insulin sensitivity

## Abstract

Angiotensin II induced increase in hypertension enhances oxidative stress and compromises insulin action and pancreatic function. Quercetin-rich foods are beneficial for hypertensive and diabetic animals owing to their antioxidant function. The aim of this study was to evaluate the antioxidant effects of quercetin in hypertensive rats on insulin action, signaling, and secretion. Wistar rats were randomly divided into three groups: sham, hypertensive rats (H), and hypertensive rats supplemented with quercetin (HQ). After three months of initial hypertension, quercetin was administered at 50 mg/kg/day for 30 days. Our results indicate that hypertension and serum lipid peroxidation levels were reduced by quercetin supplementation. We observed increased insulin sensitivity in adipose tissue, corroborating the insulin tolerance test, HOMA index, and improvements in lipid profile. Despite normal insulin secretion at 2.8 and 20 mM of glucose, animals treated with quercetin exhibited increased number of islets per section; increased protein expression of muscarinic receptor type 3, VEGF, and catalase in islets; and hepatic mRNA levels of *Ide* were normalized. In conclusion, supplementation with quercetin improved insulin action and prevented pancreatic and metabolic dysfunction.

## Introduction

Hypertension is known to be a major risk factor for heart attack, stroke, and kidney damage, affecting approximately 1 in 4 men and 1 in 5 adult women (WHO 2020b), and is also related to premature death. The World Health Organization has predicted a 25% decrease in the cases of hypertension (WHO 2020a). Renovascular hypertension is described as partial stenosis caused by renal artery obstruction (Herrmann and Textor 2019), mostly associated with atherosclerosis, decreased kidney blood flow, and activation of the renin-angiotensin system, which increases blood pressure (Basso and Terragno 2001; Faselis et al. 2011; Sabrane et al. 2005). In addition, renovascular hypertension is associated with risk factors such as diabetes, smoking, peripheral vascular disease, and coronary artery disease (Ritchie et al. 2014).

The experimental renovascular hypertension 2K1C model was described by Goldblatt, in which artery stenosis promoted by metal clips on the left renal artery produces an increase in blood pressure and stimulates the renin-angiotensin system, which produces angiotensin II (Ang II) (Goldblatt et al. 1934; Navar et al. 1998). Ang II can induce oxidative stress in the pancreatic, adipose, muscle, and cardiac tissues via NADPH (Sowers 2002, 2004; Chan and Leung 2011). Moreover, the effects of Ang II on the pancreas include morphofunctional damages, such as a decrease in insulin and blood flux in pancreatic islets (Tikellis et al. 2004; Ihoriya et al. 2014) and impaired muscle insulin signaling (Sowers 2004).

Antioxidant substances delay oxidative damage and maintain homeostasis. These substances modulate inflammation and protect against cell damage (Rahal et al. 2014; Xiao and Hogger 2015; Zhao et al. 2017). The flavonoid quercetin is involved in glycemic homeostasis, insulin sensitization, and secretion (Eid and Haddad 2017)promotes oxidative balance by modulating the activities of superoxide dismutase (SOD) and catalase (Xu et al. 2019; Msauko Koboria 2015), and acts on β cell homeostasis and oxidative protection, suggesting that quercetin is an anti-diabetic agent (Kang et al. 2019; Bule et al. 2019; Ghorbani et al. 2019; Ali et al. 2020; Bardy et al. 2013). Furthermore, studies have shown anti-hypertensive, anti-inflammatory, and anti-oxidative actions of quercetin, in addition to reducing the hypertensive response to Ang II (Marunaka et al. 2017; Salehi et al. 2020; Aprotosoaie et al. 2016).

Considering the therapeutic potential of quercetin, we evaluated the molecular and functional adaptations of the endocrine pancreas and insulin-target tissues through supplementation with quercetin in hypertensive animals. We believe that this research will improve the understanding of insulin action and pancreatic islet functionality in managing glucose homeostasis through hypertension.

## Material and Methods

### Animals

All experimental procedures were approved by the Ethics Committee of the Herminio Ometto Foundation University Center (protocol no. 013/2019) and were conducted according to the guidelines of the Brazilian College of Animal Experimentation (COBEA). Animals were treated in accordance with the Guide for the Care and Use of Laboratory Animals (8th edition, National Academies Press).

Twenty-four male Wistar rats at 40 days of age were used. Average weight was approximately 180 g and rats were kept in individual cages under a constant temperature of 22 °C with 12 h light/dark cycles and water and commercial feed ad libitum. To reproduce the renovascular hypertension model, the Goldblatt technique (Goldblatt et al. 1934) (2K1C - 2 kidneys 1 clip) was used, and the rats were anesthetized with a combination of ketamine (100 mg/kg) and xylazine (10 mg/kg) injected intraperitoneally. After an abdominal incision, the renal artery was located and dissected, and a silver clip with a 0.2 mm opening was placed over the left renal artery (H), obstructing 70% of the blood passage. The animals in the control group (sham) underwent the same procedure; however, there was no placement of the clips. Twelve weeks after surgery, the Wistar rats were separated into three groups (n = 8): sham, hypertensive (H), and hypertensive + quercetin (HQ). HQ animals received quercetin (Sigma®) supplementation of 50 mg/kg/day orally for 30 days (Figure 1A), while animals in the H and SHAM groups received vehicle orally (carboxymethyl cellulose at a concentration of 0.05%) for the same period. The body weight of the animals was assessed weekly throughout the experiment, while blood pressure was measured after the start of gavage until euthanasia. Figure 1 A shows the experimental time-line.

**Fig 1.**
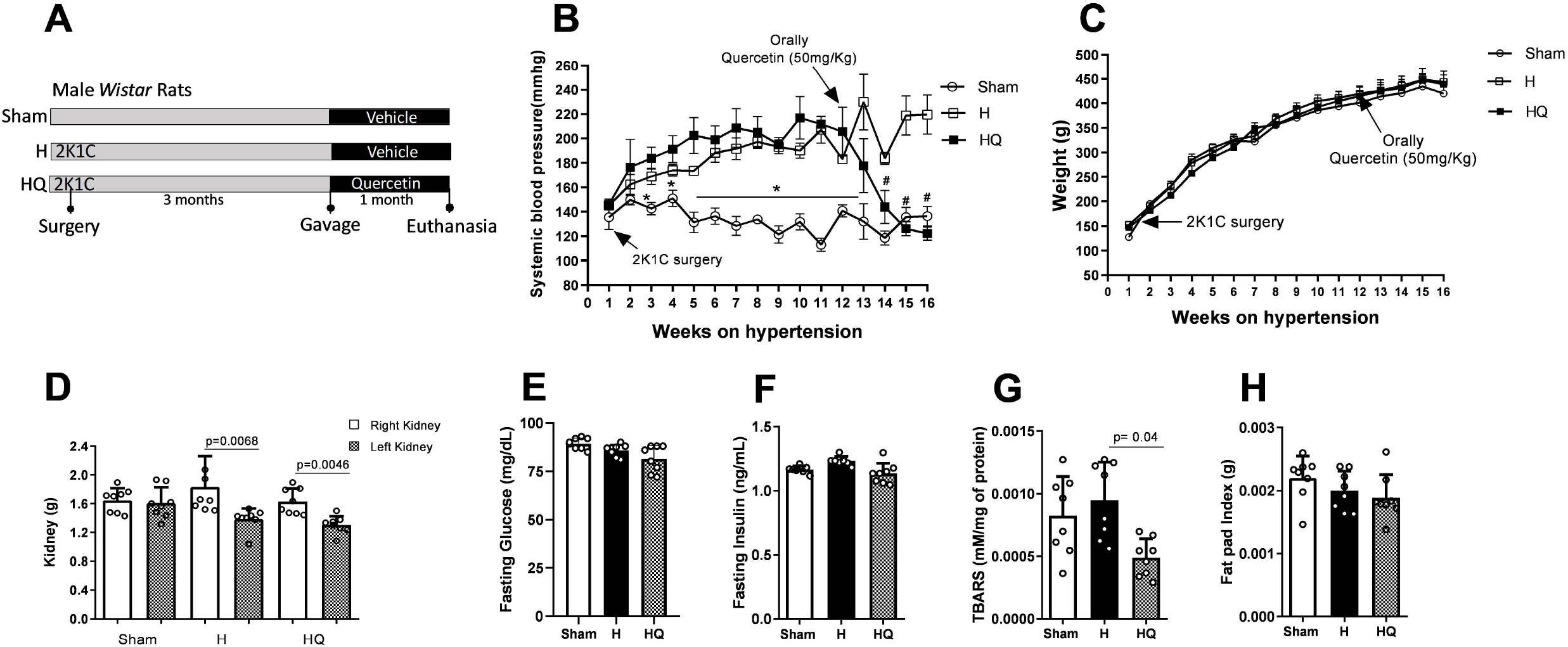
Time line (A), blood pressure (B), body weight (C), kidney weight (D), blood glucose (E), blood insulin (F), periepididymal adipose tissue index (G), and serum TBARS (H) (mean ± standard error of the mean) * one way ANOVA with Tukey Test and Student’s t-test for B, Student’s t-test for D, and ANOVA with Tukey Test for H.

### Evaluation of systolic blood pressure

Systolic blood pressure (SBP) was measured using the caudal plethysmography method (EFF 306; Insight, Ribeirão Preto-SP, Brazil). The animals were placed in a chamber heated to 37 °C. After raising the body temperature, they were placed on the plethysmograph, and a flow cuff was placed on the animals coupled to a pressure transducer. The results were calculated as the average of three consecutive measurements per animal.

### Intraperitoneal glucose tolerance test (ip.GTT) and intraperitoneal insulin tolerance test (ip.ITT)

The animals were subjected to Ip.GTT and Ip.ITT after a 6 h fast. Blood was collected from the tail of each rat (time zero). Subsequently, glucose (2 g/kg) or insulin (1.5 U/kg) were administered intraperitoneally, and blood samples were collected at each time point. The glucose concentrations in the blood samples were determined using a glucometer (OptiumXceed, Abbott). Kitt was calculated from the slope of the regression line obtained with log-transformed glucose values between 0 and 30 min after insulin administration (Bonora et al. 1989). Homeostasis model assessment of insulin resistance (HOMA-IR), homeostasis model assessment of insulin sensitivity (HOMA%S), and homeostasis model assessment of β-cell function (HOMA-β) were calculated using the website www.dtu.ox.ac.uk/homacalculator.

### Hormonal and biochemical assessments

Biochemical analyses of glucose, triglycerides, cholesterol, LDL, and HDL were performed using commercial kits (Laborlab, Guarulhos, SP, Brazil). Insulin was measured by ELISA (EZRMI-13K Sigma-Aldrich - Rat/Mouse - Insulin ELISA) and the hepatic and skeletal muscle glycogen contents of fed rats (Lo et al. 1970) were measured by a colorimetric method. This method is based on the degradation of glycogen to glucose by the phenol-sulfuric acid and the absorbance of the characteristic yellow-orange color was measured at 490 nm.

### Serum tiobarbituric acid reactive substances (TBARS)

Determination of TBARS was performed as described previously (Esterbauer and Cheeseman 1990), by measuring malondialdehyde (MDA), a product of lipoperoxidation primarily formed due to reaction with hydroxyl free radicals. The TBARS concentration was determined spectrophotometrically at 532 nm and expressed as nmol of MDA per mg of protein.

### Fat pad tissue and kidney weight index (g)

Weights of the right and left kidney, periepididymal adipose tissue, and total body were measured. The fat pad index was calculated by dividing the weight of the adipose tissue by the total body weight.

### Secretion by isolated islets

Islets were isolated manually after collagenase digestion of the pancreas. Groups of five islets were first incubated for 30 min at 37 °C in Krebs-bicarbonate solution containing 5.6 mmol/L glucose and equilibrated with 95% O_2_/5% CO_2_, pH 7.4. The solution was then replaced with fresh Krebs-bicarbonate buffer, and the islets were incubated for an additional hour in the presence of glucose (2.8 and 20 mmol/L). The incubation medium contained: 115 mmol/L NaCl, 5 mmol/L KCl, 10 mmol/L NaHCO_3_, 2.56 mmol/L CaCl_2_, 1 mmol/L MgCl_2_, 15 mmol/L HEPES, and 0.3% BSA (w/v). The insulin content of each sample was measured using an ELISA kit (Biotec Center, SPI-Bio).

### Gene expression by RT-PCR

Total RNA was isolated from approximately 100 mg rat livers using TRIzol® reagent (Invitrogen, Carlsbad, CA, USA), followed by the digestion of contaminating DNA with DNAse I (amplification grade; Invitrogen), according to the manufacturer’s instructions. RNA purity and concentration were determined spectrophotometrically. cDNA was synthesized from 2 μg RNA using High-Capacity cDNA Reverse Transcription (Applied Biosystems™) kit in a final volume of 20 μL. The mRNA levels of insulin degrading enzyme (*Ide*), acetyl-CoA carboxylase (*Acc1*), and 3-hidroxi-3-methyl-glutaril-CoA reductase (*Hmgco-A*) genes were investigated by semi-quantitative RT-PCR. The primer sequences used in the PCR reactions were chosen based on sequences available in GenBank. *Ide* was amplified using gene-specific forward 5’-AGGAAATGTTGGCTGTGGACGCA-3′ and reverse 5′-CCTGGCAAGAACGTGGACGGATA-3’ primers which produced a predicted amplicon of 62 bp (Tm 57 °C). *Acc1* was amplified using forward: 5′TGGATGAACCATCTCCGTTG3′ and reverse: 5-‘GTTTTACTAGGTGCAAGCCG3′ primers and produced a predicted amplicon of 255 bp (Tm 57 °C); and *Hmgco-A* was amplified using forward: 5′CTTGACGCTCTGGTGGAATG3′ and reverse: 5′TGGCTGAGCTGCCAAATTGG3′ primers which amplified a predicted amplicon of 183 bp (Tm 57 °C). Amplified products were separated on a 2.0 % agarose gel and stained with ethidium bromide. The gel was photographed using Syngen G: Box. Signal intensities of the bands were measured densitometrically using the ImageJ software. Each value was determined as the mean of three densitometric readings. Results were expressed as average ratios of the relative expression of transcripts normalized with β*-actin* as the control housekeeping gene (forward: 5′AGAGGGAAATCGTGCGTGACA3′ and reverse: 5-′CGATAGTGATGACCTGACCGTCA3), producing a predicted amplicon of 138 bp (Tm 57 °C).

### Western blotting

After a 12-to 14-h fasting period, the rats were anesthetized by intraperitoneal injection of ketamine and xylazine and opened 10–15 min later, i.e., as soon as the effect of anesthesia was ensured by the loss of pedal and corneal reflexes. The abdominal cavity was opened, the portal vein was exposed, and 0.2 mL normal saline was injected with or without insulin (10^−6^ M). At 15 min after insulin injection, the liver and epididymal adipose tissues were removed and frozen. Isolated pancreatic islets were subjected to western blotting. The protein concentration was determined using the biuret method. Aliquots of the supernatant were treated with Laemmli buffer. Proteins (50 μg) were loaded onto 12–15% SDS-PAGE gels using a Mini-Protean apparatus (Bio-Rad, CA, USA) and transferred onto nitrocellulose membranes (Bio-Rad). The membranes were incubated in blocking solution (5% BSA), after which the membranes were incubated at 4 °C overnight with polyclonal antibodies. Antibodies against catalase (sc-34284), SOD-2 (sc-137254), PCNA (pc-10), mAchM1 (c-20), mAchM3 (H-210), VEGF (sc-53462), Insulin Rβ (sc-711), p-Insulin Rβ (sc-25103), β-actin (sc-47778) were purchased from Santa Cruz Biotechnology (CA, USA;1:200). Antibodies against AMPK (2532), p-AMPK (2535), AKT (9272), and p-AKT (9271) were purchased from Cell Signaling (1:1000). This was followed by incubation with secondary antibodies (Santa Cruz Biotechnology, CA, USA) at a dilution of 1:10000 and exposure to the SuperSignal West Pico Chemiluminescent Substrate kit. The band intensities were quantified by optical densitometry using the free Scion Image (Scion Corporation) software. The β-actin (1:5000, Santa Cruz Biotechnology, CA, USA) was used as an internal control.

### Histological analysis of pancreas

After fixation and dehydration of the pancreas, small fragments were embedded in paraffin using standard procedures. Sections (5 μm thick) were stained with hematoxylin-eosin solution and images were acquired using a Leica DM 2000 photomicroscope and LAS v.4.1 software. For each animal, 40 images were obtained at 400× magnification (1032 × 1040 pixels). ImageJ software was used to quantify the area of each islet in each image. Furthermore, the islet numbers per section were determined.

### Immunohistochemistry

For the immunohistochemical study, antigens were recovered from 5 μm thick sections using sodium citrate buffer (10 mM, pH 6.0) at 100 °C for 40 min. Endogenous peroxidases and non-specific binding were blocked using a Novolink Max Polymer Detection System (RE7280-K; Leica Biosystems®). The samples were incubated with the primary antibody (PCNA – pc10 1:200; Santa Cruz Biotechnology) overnight. Subsequently, the tissues were incubated with Novolink Polymer (Leica Biosystems), the reaction was developed with diaminobenzidine, and the sections were counterstained with Harris hematoxylin. Five images from each animal at 400× magnification were captured using a Leica DM2000 microscope, and quantified using ImageJ v.1.53e (NIH, USA).

### Statistical analysis

Data are shown as the mean ± standard error of the mean. All data were analyzed using the Shapiro-Wilk Normality test. For the data that fit the normality curve, the parametric Student’s t-test was used to compare two groups at different experimental times. For comparison between the different experimental times only in the HQ group, one-way ANOVA and Tukey post-test were used. All statistical and graphical tests were performed using GraphPad Prism software (version 8.0). The pre-established significance level was set at P <0.05.

## Results

### Quercetin supplementation decreases arterial pressure and TBARS levels in 2K1C rats

Quercetin reduced the BP (blood pressure) of HQ animals to that observed in sham animals (Figure 1B). The decrease in BP in quercetin animals was observed from the 13th week. At the end of the 16th week, the blood pressure of the animals in the HQ group was equal to that of the animals in Sham group. There were no significant differences in the weight of the animals (Figure 1C). Subsequently, we compared the hypertrophy in right kidney of the animals in H and HQ groups with that of the left kidney, verifying and validating the animal model of renovascular hypertension (Figure 1D). Blood glucose (Figure 1E), insulinemia (Figure 1F), and adipose tissue (Figure 1H) were similar in both groups. TBARS serum was lower in the HQ group than in the other groups, which is consistent with the antioxidant effect of quercetin (Figure 1G).

### Quercetin-modulated serological and hepatic lipid metabolism in 2K1C animals

To investigate whether quercetin induced changes in the lipid profile, we evaluated the serological profile of lipids and key enzymes involved in lipid synthesis. Figure 2 illustrates the values obtained from biochemical analyses of the serum and liver, as well as gene expression in the liver tissue. Animals in the sham, H, and HQ groups showed similar concentrations of cholesterol (Figure 2B), HDL (Figure 2C), hepatic triglycerides (Figure 2E), hepatic cholesterol (Figure 2F), and hepatic glycogen (Figure 2G). In addition, a previous report showed a decrease in triglyceride (Figure 2A) and LDL (Figure 2D) levels in quercetin-animals (Ikewuchi et al. 2013), corroborating our data. *Acc1* mRNA levels (Figure 2H), key enzymes for fatty acid biosynthesis, and *Ide* (Figure 2J), which is responsible for insulin clearance, were normalized in the quercetin-supplemented group to a level similar to that of the Sham group. We did not observe any statistical differences in *Hmgco-A* (Figure 2I), which is responsible for cholesterol synthesis. In Figure 3K, the representative bands of RT-PCR referring to Figures 2 H, I, and J can be seen.

**Fig 2.**
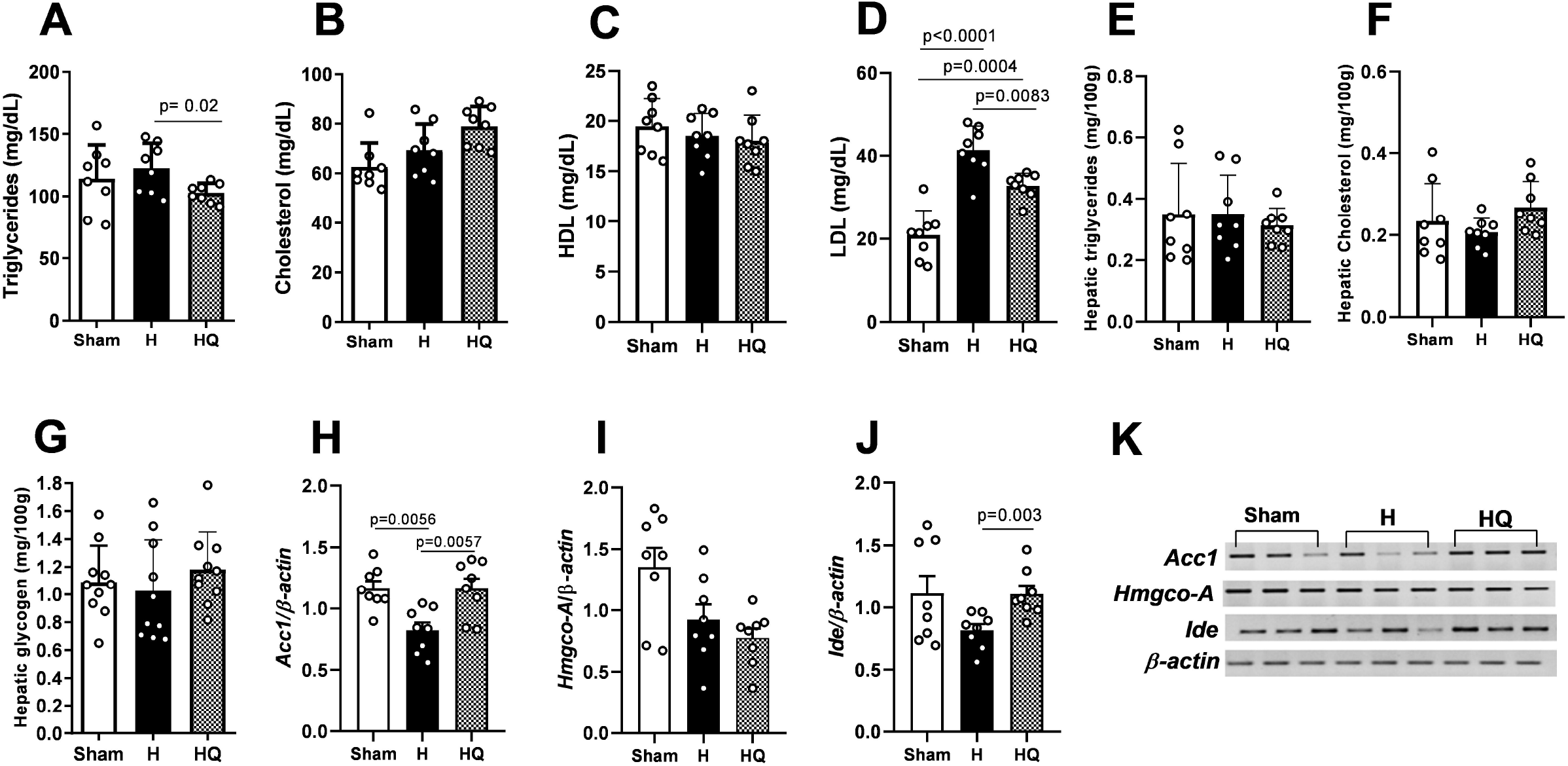
Serum triglycerides (A), cholesterol (B), HDL (C), LDL (D), hepatic triglycerides (E), hepatic cholesterol (F), hepatic glycogen (G), *Acc1* gene (H), *Hmgco-A* (I) and *Ide* (J), representative panel of the RT-PCR bands of the genes (K) (mean ± standard error of the mean, one-way ANOVA followed by post hoc Tukey).

**Fig 3.**
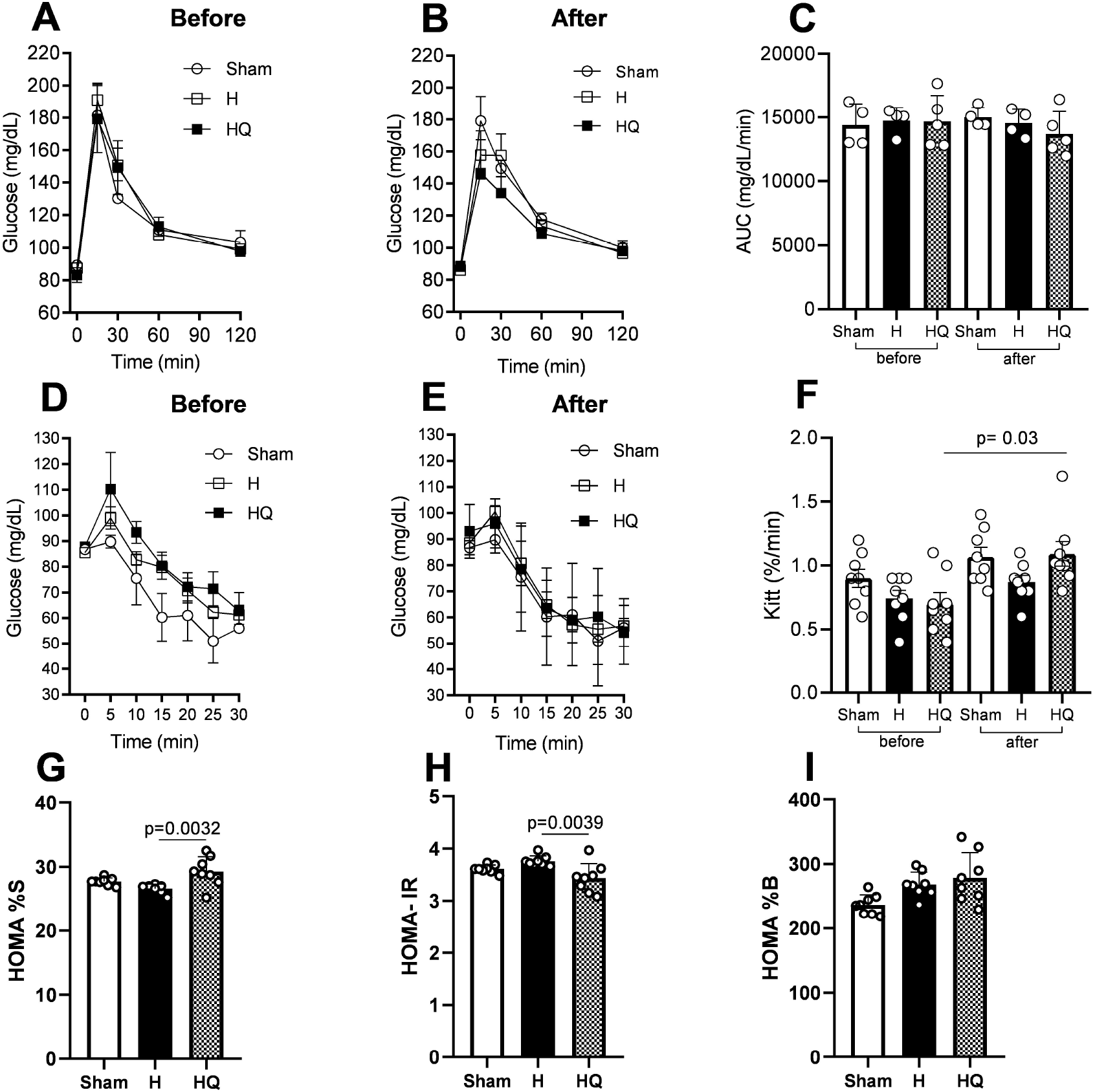
(A) Intraperitoneal Glucose Tolerance Test (ip.GTT) before supplementation with quercetin, (B) GTT after supplementation, (C) area under the curve (AUC), (D) Intraperitoneal Insulin Tolerance Test ipITT before supplementation, (E) ITT after supplementation, (F) glucose disappearance constant (Kitt), HOMA%S (G), HOMA-IR (H), HOMA%B (I) (mean ± standard error of the mean, one-way ANOVA followed by post hoc Tukey).

### Increased insulin sensitivity and glycemic homeostasis in 2K1C animals supplemented with quercetin

To investigate glucose tolerance and insulin sensitivity, we performed GTT and ITT. In addition, we investigated the association between insulin resistance and renovascular hypertension using a mathematical model of the HOMA. Figure 3 shows the GTT and ITT performed before and after supplementation with quercetin. Test analyses showed similar results and suggested glycemic homeostasis in all animal groups (A, B, C, D, and E). However, the glucose decay rate (Figure 3F) during ITT was higher for HQ animals, suggesting an increase in insulin sensitivity compared to animals in the same group before supplementation with quercetin. The HOMA model estimates beta-cell function (%B) and insulin sensitivity (%S). HOMA%S (Figure 3G) was higher in the HQ group than in the H group, while the HOMA-IR (insulin resistance) was lower in the HQ group than in the H group (Figure 3H), suggesting increased insulin sensitivity, which can be attributed to quercetin. The HOMA%B index (Figure 3I) was slightly higher in the HQ group than in the H group (p = 0.726) and was statistically higher than that in the sham group, suggesting a beneficial effect on pancreatic islet homeostasis.

### Increased tissue-specific insulin sensitivity in animals treated with quercetin

We investigated the effects of quercetin supplementation in 2K1C animals at the initiation of insulin signaling in key target tissues. Interestingly, in the liver, we did not observe any significant differences in the levels of IR, AKT, and AMPK phosphorylation (Figure 4A, 4B, and 4C). However, the levels of IR and AKT phosphorylation in adipose tissue increased in HQ animals after insulin stimulation (Figure 4F and 4G). Additionally, we also found an increase in AMPK protein levels in the HQ group compared to that in the H group. As expected, in the liver and adipose tissue of the HQ group, the protein expression levels of the antioxidant proteins SOD-2 (Figure 4D, 4I) and catalase (Figure 4E and 4J) were higher than those in group H.

**Fig 4.**
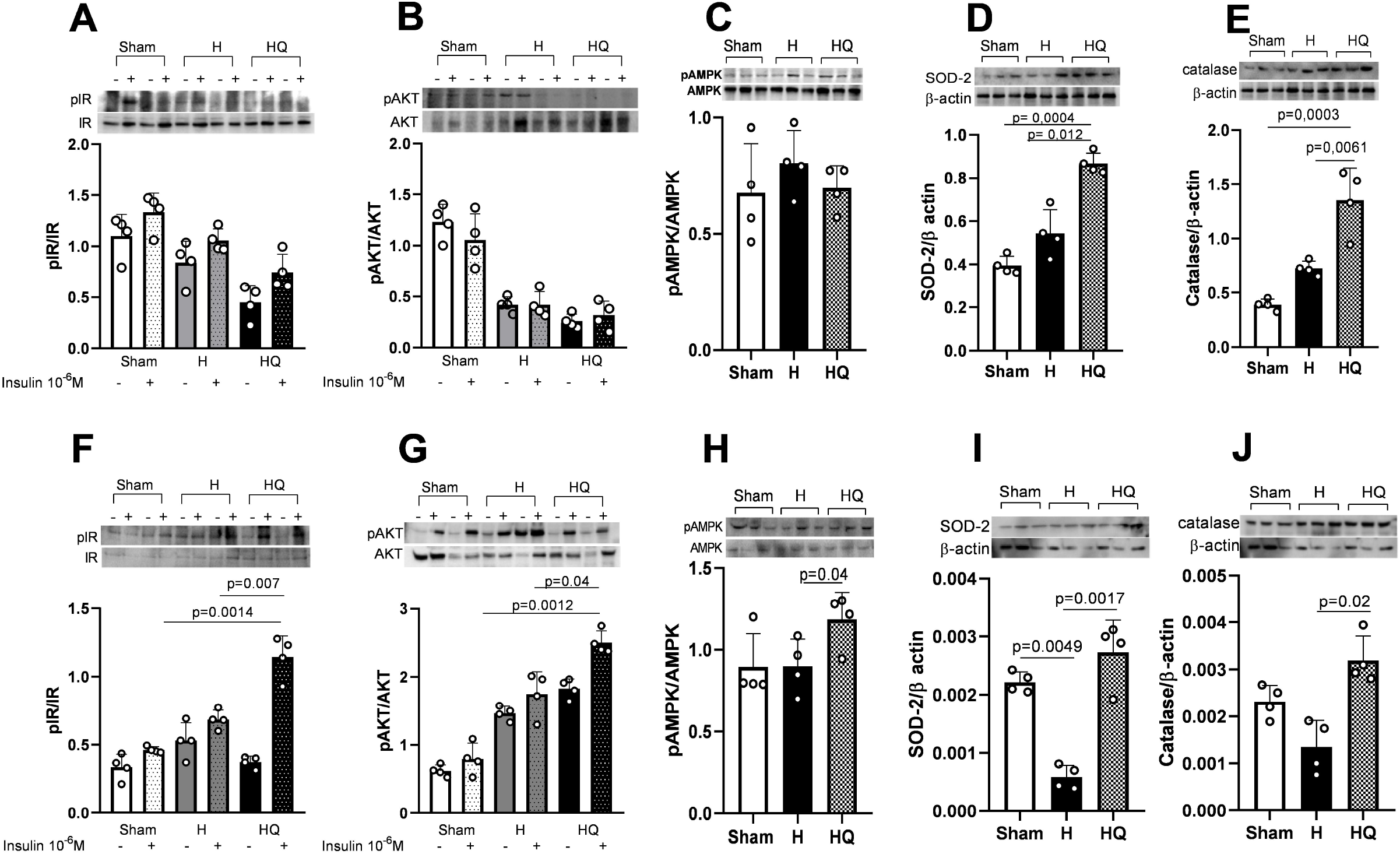
In liver tissue, we observed IR (insulin substrate) phosphorylation (A), AKT phosphorylation (B), AMPK phosphorylation (C), SOD-2 (D), and catalase (E). In adipose tissue we observed phosphorylation of IR (insulin substrate) (F), the phosphorylation of AKT (G), the phosphorylation of AMPK (H), SOD-2 (I), and catalase (J) (mean ± standard error of the mean, one-way ANOVA followed by post hoc Tukey).

### Quercetin promotes increased protein expression of the muscarinic M3 receptor, catalase, and VEGF in isolated islets

We evaluated whether the metabolic improvements induced by quercetin were accompanied by molecular and functional changes in the pancreatic islets. Figure 5 illustrates the values obtained from protein and insulin secretion analyses of isolated pancreatic islets associated with morphometric and immunohistochemical analyses performed on pancreatic paraffin sections. Protein expression of M3 (Figure 5B), catalase (Figure 5E), and VEGF (Figure 5F) receptors in isolated islets increased in the animals of the HQ group compared to that in the animals of the H group. Similar results were found for M1 muscarinic receptors (Figure 5A) and PCNA (Figure 5A). 5G). However, we did not observe an increase in SOD-2 in H versus HQ animals (Figure 5C). Figure 4H illustrates insulin secretion under sub-physiological (2.8 mM) and supraphysiological (20 mM) glucose stimulation. We did not observe differences in insulin secretion stimulated by low and high glucose concentrations in all groups. As shown in Figure 5I, we observed a greater number of islets per section in HQ animals than in H animals, suggesting that quercetin plays a role in the maintenance of islets. In figures 5J and 5K, we observed similarities in islet area and number of PCNA-positive cells. In figure 5D, we present the representative bands of western blotting and in figure 5L the illustrative images of islets stained by HE and immunohistochemistry for PCNA are shown.

**Fig 5.**
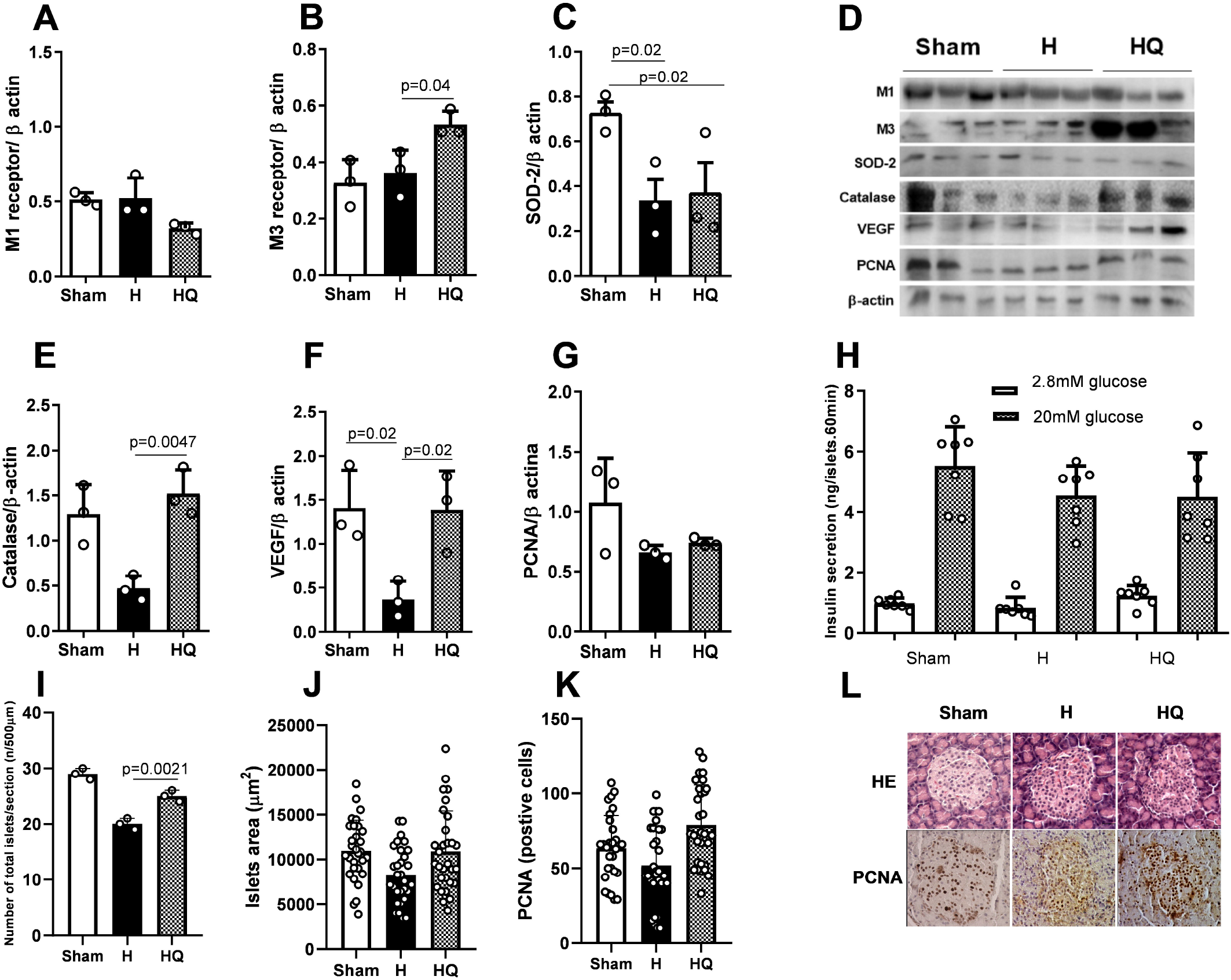
Protein expression of M1 (A), M3 (B), SOD-2 (C), representative panel of western blotting bands (D), protein expression of catalase (E), VEGF (F), PCNA (G), secretion of insulin per isolated islets stimulated in 2 mM and 20 mM of glucose per animal group (H). In the paraffinized pancreatic tissues we observed the percentage of islets per section (I), islet area (J), protein expression of PCNA by immunohistochemistry and a representative panel of islets stained by HE and the representation of positive marking for PCNA (L). Specifically, for the blotting in isolated islets, the experiments were performed in triplicate (pool of islets with 200 islets per replicate). Insulin secretion was performed with n=7 (pool of three animals) (mean ± standard error of the mean, one-way ANOVA followed by post hoc Tukey).

## Discussion

Oral quercetin supplementation in hypertensive rats showed an antioxidant effect and reduced blood pressure, in addition to promoting the action of insulin and preventing tissue deterioration in pancreatic islets by ensuring cholinergic signaling and oxygen supply.

The reduction in blood pressure of hypertensive animals after supplementation with quercetin has been reported by other researchers in human studies. The literature is rich in data on overweight and obese humans, with or without diabetes, showing a reduction in blood pressure due to supplementation with quercetin at different doses and duration of treatment (Brull et al. 2015; Serban et al. 2016; Gormaz et al. 2015; Edwards et al. 2007). In overweight patients with a high risk of cardiovascular disease, administration of quercetin at a dose of 150 mg/day for six weeks reduced systolic blood pressure, oxidized LDL, and TNF-α (Egert et al. 2009). Consistent with the literature, our results show a reduction in serum triglycerides and LDL in hypertensive animals supplemented with quercetin, although the gene expression of *Hmg-coa* was similar among the groups. In this context, normalization of the levels of *Acc1* enzyme in the hypertensive animal group supplemented with quercetin, to levels similar to those in the Sham group suggest that quercetin helps in attaining normal rates of adipogenesis and fatty acid oxidation. These lipid metabolism improvements are associated with insulin sensitivity and glucose tolerance, as reported in the literature (Fullerton et al. 2013) considering that insulin regulates the *Acc1* promoter gene (Kim 1997).

Consistent with this, the HOMA%S, and HOMA-IR indices were favorable for increasing insulin sensitivity in 2K1C hypertensive animals supplemented with quercetin as well as Kitt. These adaptations led us to suggest that quercetin has anti-hyperglycemic and anti-diabetic effects through improvements in insulin sensitivity (Brull et al. 2015; Gormaz et al. 2015; Edwards et al. 2007; Serban et al. 2016), although we did not find differences in fasting glucose and insulin levels. Similar results have been reported for the HOMA index in streptozotocin-induced and quercetin-treated diabetic animal models (Ali et al. 2020).

In agreement with the insulin sensitivity data in hypertensive animals being attributed to quercetin action, we found an increase in IR and AKT phosphorylation in adipose tissue. The antioxidant action of quercetin in adipose tissue is mediated by catalase and SOD-2 upregulation. Insulin resistance and chronic hyperglycemia in type 2 diabetes are often associated with hypertension and an increased risk of cardiovascular diseases. The effect of Ang II is related to the inhibition of (phosphatidylinositol 3 kinase) PI3K and consequently AKT signaling. Thus, Ang II opposes the action of insulin in increasing glucose uptake in skeletal and adipose muscles, leading to insulin resistance (Jing et al. 2013). Quercetin, at a 10 nM dose in muscle cell culture, activated the phosphorylation of IRS1 (insulin receptor substrate 1), PI3K, and AKT, suggesting that quercetin acted in the same way as insulin, via promoting insulin AKT’s role in regulating glucose levels (Jiang et al. 2019). Quercetin also increases adipose tissue, IR phosphorylation, and consequently GLUT-4 translocation in diabetic animals, modulating insulin sensitivity (Ali et al. 2020). Specifically, in adipose tissue, the AMPK elevation induced by quercetin may suggest an association with the regulation of lipid metabolism. A previous report shows reduced hepatic lipogenesis and reduced lipid accumulation by AMPK activation, and consequently inhibition of ACC1 and ACC2 (Koistinen et al. 2003). This lipid-lowering effect decreases obesity-induced insulin resistance (Fullerton et al. 2013). These results were also reported by other researchers, who observed the activation of AMPK by quercetin in muscle cells (Dhanya et al. 2017). In the present study, we did not observe the same effects in the initial stages of the insulin signaling cascade in hepatic tissue, such as IR, AKT, and AMPK activation; however, quercetin promoted the antioxidant effect, as evidenced by catalase and SOD-2 protein expression. Corroborating these results of non-activation of the insulin cascade, we did not observe an increase in liver glycogen stores in hypertensive animals treated with quercetin. Different assessments that can measure glucose uptake by the liver under these conditions would be necessary to strengthen the discussion, but were beyond the scope of this study.

Many polyphenolic compounds have been reported to play an important role in maintaining glucose homeostasis and have a protective effect on pancreatic islets (Szkudelska et al. 2019; Kang et al. 2019). Pleiotropic quercetin mechanisms involve intestinal inhibition of glucose absorption, increased insulin secretion and sensitization, and improved glucose utilization in peripheral tissues (Eid and Haddad 2017). Cholinergic mechanisms are important for the function and survival of the endocrine pancreas (Gilon and Henquin 2001; Gautam et al. 2010), and muscarinic receptors stimulate insulin secretion and islet proliferation during pancreatic regeneration (Moulle et al. 2019; Ito et al. 2019). The literature indicates that M1 and M3 receptors are predominant in pancreatic islets and regulate insulin secretion (Fernandez-Cabezudo et al. 2019). Consistent with increase in M3, hepatic *Ide* mRNA normalization is consistent with the clearance function of insulin, amylin, and glucagon, three hormones that are vital for glucose homeostasis (Najjar and Perdomo 2019). Impairments in IDE function are associated with glucose intolerance and type 2 diabetes (Tang 2016). IDE-null mutant mice, IDE-knockout, and pharmacological inhibition of IDE in rodents resulted in hyperinsulinemia (Maianti et al. 2014), as well as glucose intolerance associated with hyperinsulinemia and insulin resistance (Abdul-Hay et al. 2011).

Associated with the cholinergic mechanism of insulin secretion, there is evidence that insulin exocytosis deficiency is caused by abnormalities in the vascular network of the islets (Brissova et al. 2014). The lower insulin secretion can be attributed to lower blood vessel density and size, which likely reduces blood flow to the islets and consequently causes difficulties in islet insulin exocytosis (Brissova et al. 2014). Although we did not observe differences in insulin secretion in pancreatic islets, we found an increase in VEGF and catalase in isolated islets of HQ animals. Studies have shown that treatment with quercetin increased cell viability and functionality of β cells by reducing H2O2 and increasing insulin secretion stimulated by glibenclamide and glucose via ERK1/2 phosphorylation in INS-1 cells (Youl et al. 2010), and inhibition of p38/MAPK phosphorylation (Youl et al. 2014). Other studies have shown that quercetin reduces hyperglycemia and protects β cells directly through the elimination of lipid peroxides and increased production of antioxidant enzymes (Lacraz et al. 2009), corroborating our TBARS serological findings. Thus, the increase in VEGF observed in islets of hypertensive animals supplemented with quercetin suggests an association with the preservation of insulin secretion and maintenance of the number of pancreatic islets. Confirming our view, the literature showed that Ang II is involved in the reduction of glucose-stimulated insulin secretion and the regulation of pancreatic microcirculation (Ihoriya et al. 2014; Ghorbani et al. 2019). Although the levels of PCNA, a cell proliferation marker, were similar between groups, we observed a restoration of the number of islets after supplementation with quercetin. Furthermore, the increase in M3 receptors may be associated with the normalization of the number of islets, as previous studies have shown that pancreatic islet proliferation occurs via activation of the parasympathetic nervous system via the vagus nerve (Moulle et al. 2019). Quercetin injection at a dose of 15 mg/kg for 10 days promotes regeneration of pancreatic islets by inducing an increase in the number of pancreatic islets and consequently increased insulin secretion, which is reflected in a reduction in plasma glucose, cholesterol, and triglycerides (Vessal et al. 2003). Other studies have shown that quercetin potentiates insulin secretion in INS-1 cells (Youl et al. 2010) and INS-1E cells (Bhattacharya et al. 2014). These effects suggest the mechanisms of transient inhibition of KATP channels and transient activation of voltage-dependent Ca2+ channels (Kittl et al. 2016).

## Conclusion

In conclusion, our results indicate that quercetin reverses the increase in blood pressure, providing an improved clinical condition in an animal model. This was evidenced by the improvements in the lipid profile and increase in insulin sensitivity through the activation of the initial steps of the insulin signaling cascade in adipose tissue, followed by a protective mechanism against the action and secretion of pancreatic islets.

## Author contributions

All authors performed all the experiments. B. Calsa and Maria Esméria Corezola do Amaral analyzed the results and revised the manuscript; Maria Esméria Corezola do Amaral designed the study and wrote the manuscript.

## Acknowledgments

The authors are grateful to Bruno Alves Cia, BSc, Sr Mateus Eduardo Bortolanza da Silva, Sra Ana Cristina Pires Menegheti, Renata Barbieri, and Lucas Eduardo Orzari for their excellent technical assistance.

